# Nuclear export in somatic cyst cells controls cyst cell-germline coordination and germline differentiation in the *Drosophila* testis

**DOI:** 10.1101/452466

**Authors:** Fani Papagiannouli, Margaret T. Fuller, Ingrid Lohmann

**Affiliations:** Institute for Genetics, University of Cologne, Cologne, 50674, Germany; Department of Developmental Biology, Beckman Center, Stanford University School of Medicine, CA 94305-5329, Stanford, USA; Centre for Organismal Studies (COS) Heidelberg, Cell Networks – Cluster of Excellence, University of Heidelberg, 69120, Heidelberg, Germany

**Keywords:** *Drosophila* testis, germline, soma-germline communication, spermatogenesis, nucleoporin, exportin, profilin, Rae1, Nxt1, Ntf-2, nuclear export

## Abstract

Nucleocytoplasmic communication is crucial for proper cell function and coordination of intrinsic cues with signaling responses emanating from the neighboring cells and the local tissue microenvironment. In the *Drosophila* male germline system, germ cells proliferate and progressively differentiate enclosed in supportive somatic cyst cells, forming a small cyst, the functional unit of differentiation. Here we show that the peripheral nucleoporins Nup62, Nup214 and Nup88, and the exportin Emb are critically required in cyst cells to maintain cyst cell survival and germline encapsulation in order to protect cyst cell-germline communication and promote germ cell differentiation. Knockdown of *nup62, emb, nup214* or *nup88* in cyst cells leads to cell-autonomous defects in mRNA export, and cell non-autonomous overproliferation of early germ cells in the absence of cyst cell-derived differentiation signals. Suppression of apoptosis can reverse cyst cell elimination and partially restored those defects. Interestingly, overexpression of the *Drosophila* Profilin gene *chickadee* can rescue cyst cell survival and restore germline encapsulation and differentiation, by counteracting Ntf-2 mediated export, suggesting that the function of Profilin in cyst cells is linked to nuclear export.

## INTRODUCTION

Adult stem cells are the lifetime source of many types of differentiated cells. They reside in microenvironments, the stem cells niches, a term first introduced in 1978 by Ray Schofield, who described the physiologically limited microenvironment that supports the hematopoietic stem cells (HSCs) (Schofield, 1978). Stem cell niches have critical roles as organizers that control stem cell maintenance and balance self-renewal vs. differentiation. Tissue-specific stem cells are regulated through their own intrinsic program as well as by signals emanating from their microenvironment. Tissues rely on stem cells to constantly replenish their cellular populations under normal conditions but also activate proliferation of resident stem cell upon damage, injury or aging to facilitate tissue regeneration (Fuller and Spradling, 2007; Matunis et al., 2012; Papagiannouli and Lohmann, 2015). Therefore, stem cell proliferation needs to be tightly regulated, since on one side stem cells must be able to respond appropriately to grow demands while at the same time they should limit the potential risk of aberrant proliferation and tumorigenesis.

The *Drosophila* testis contains two types of stem cells: the GSCs and the somatic cyst stem cells (CySCs). Each GSC is flanked by two somatic cyst stem cells (CySCs) and both types of stem cells are maintained through their association to the hub cells (Fig. 1A). Upon asymmetric cell division, each GSC produces a new GSC attached to the hub and a distally located gonialblast (GB). The CySCs also divide asymmetrically to generate a CySC remaining associated with the hub and a distally located post-mitotic daughter somatic cyst cell (CC) (Fuller and Spradling, 2007). The gonialblast divides mitotically four more times, in so-called transit-amplifying divisions (TA), giving rise to 2-, 4-, 8- and eventually 16- interconnected spermatogonial cells. After completion of TA divisions, spermatogonia become spermatocytes, undergo pre-meiotic DNA replication that turns on the transcriptional program for terminal differentiation, and undergo meiosis. Cyst cells co-differentiate with the germ cells they enclose, grow enormously in size, elongate and encapsulate them throughout their differentiation steps up to individualization and sperm production in the adult testis (Fuller, 1993; Papagiannouli and Lohmann, 2015).

**Figure 1.**
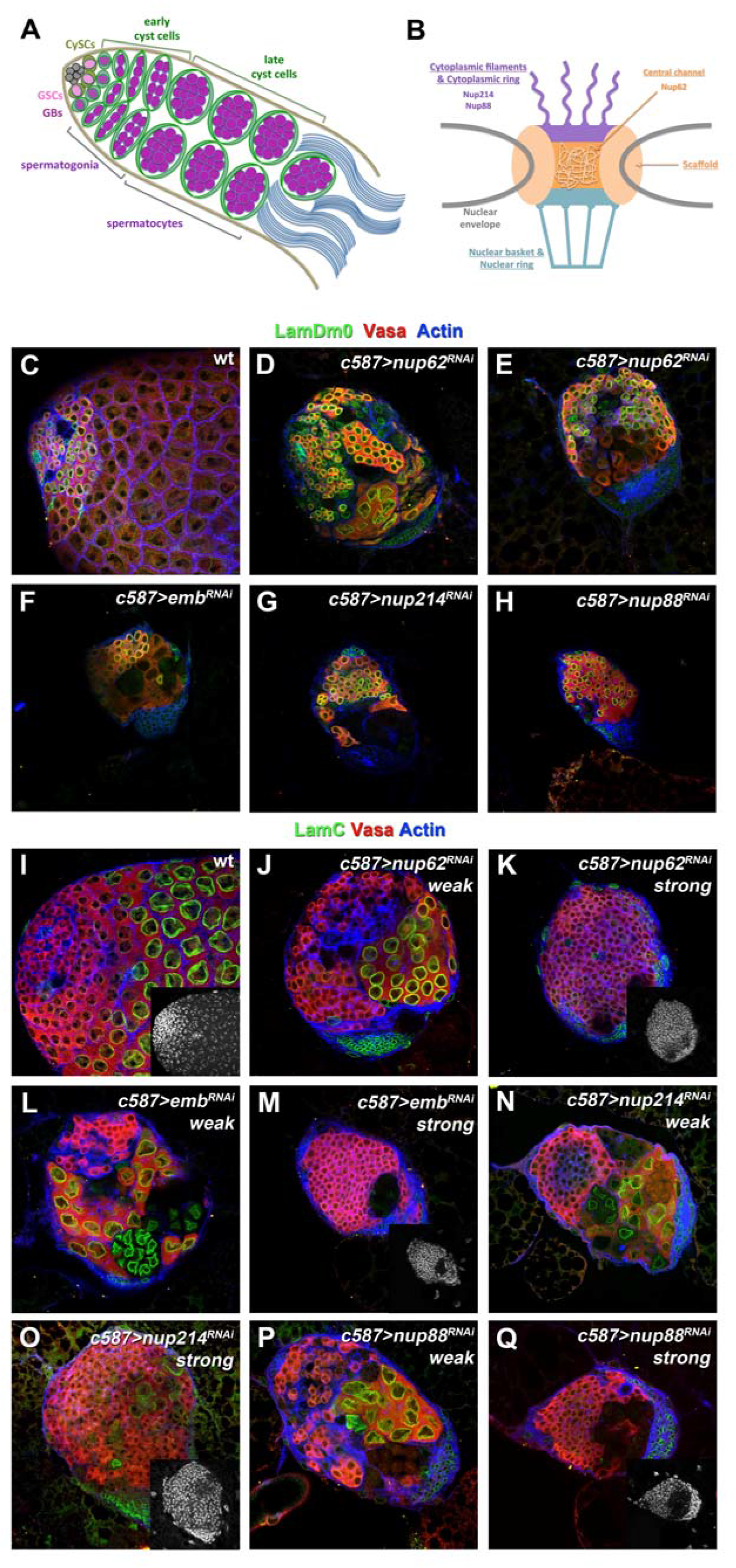
**Nup62, Nup214, Nup88 and exportin Emb have critical functions in cyst cells associated with cell non-autonomous effects in the germline. (A)** Diagram depicting early *Drosophila* spermatogenesis. GSC: germline stem cell, CySC: somatic cyst stem cell, GB: gonialblast. **(B)** Schematic diagram of the nuclear pore structure depicting the localization of the nucleoporin proteins analyzed in this study. **(C-Q)** Stainings of L3 testes of the indicated genotype for Vasa (red) marking the germline, Actin (blue) with phalloidin marking the cyst cells, the germline spectrosome and fusome, and (C-H) LamDm0 (green) marking cyst cells and early germ cells (GSCs, GBs and spermatogonia) or (I-Q) LamC (green) marking the spermatocytes. Small inset pictures (I, K, M, O, Q) show DAPI-bright germ cells. Testes are oriented anterior left. Image frame: 246μm This figure is associated with **Figure S1**.

Gamete development and mature sperm production requires a coordinated soma-germline interaction that keeps stem cell maintenance of GSCs and CySCs around the hub but at the same time allows their progenies to enter the differentiation program, which is maintained by cyst cell-germline communication away from the niche. Cyst cells always as a pair enclose the germ cells creating a cyst microenvironment that maintains cyst integrity and supports testis differentiation and morphogenesis (Zoller and Schulz, 2012). Cyst cells provide an environment necessary to trigger germline differentiation, as early germ cells with GSC characteristics can be maintained over time after ablation of all CySCs and daughter cyst cells (Lim and Fuller, 2012). In particular, driving the pro-apoptotic gene *grim* in cyst cells eliminates them and results in germ cells that proliferate massively away from the hub, fail to enter the TA mitotic program and never become spermatocytes. Thus, soma-germline coordination shapes spermatogenesis from stem cell maintenance and differentiation up to mature sperm production (Fabian and Brill, 2012; Leatherman, 2013; Zoller and Schulz, 2012).

The massive overproliferation of the germline, resembling early-stage germ cell tumors in the absence of germline differentiation, is not only manifested when cyst cells are dying (Lim and Fuller, 2012). It also appears, as a cell non-autonomous effect, whenever cyst cells are unable to encapsulate properly the germline or whenever signals normally emanating from cyst cells are lost, resulting in impaired germline differentiation (Ayyub et al., 2015; Dominado et al., 2016; Gonzalez et al., 2015; Li et al., 2003; Qian et al., 2015; Saito et al., 2009; Shields et al., 2014; Tamirisa et al., 2018). Knockdown of the *Drosophila* homologue of Profilin, encoded by the *chickadee (chic)* gene, in the cyst cells impairs germ cell enclosure, restricts TA spermatogonial divisions and results in early germ cell overproliferation, while cyst cells are progressively lost over time (Shields et al., 2014).

Proper communication between the nucleus and the cytoplasm is critically important for a coordinated cell-cell communication via signaling pathways and the intersection with intrinsic cues, since cellular responses often relay on shuttling of various components in and out of the nucleus. At the center of nucleocytoplasmin transport and communication is the nuclear pore complex (NPC), a multiprotein channel that spans the double-lipid bilayer of the nuclear envelope (Raices and D’Angelo, 2012). The NPC allows selective nucleocytoplasmic transport of macromolecules, primarily through the function of exportins and importins (collectively called karyopherin proteins) that facilitate the transport of proteins and RNAs outside and inside of the nucleus, respectively. NPCs consist of multiple copies of roughly 30 different proteins known as nucleoporins (NUPs) that form the scaffold embedded in the nuclear membrane and consists of: a central channel made of phenylalanine-glycine (FG)-containing NUPs; a cytoplasmic ring with cytoplasmic filaments facing the cytoplasm; and the nuclear ring and nuclear basket facing the nucleoplasm (Fig. 1B). The protein composition of NUPs varies among different cell types and tissues, and in support of that, mutations in various NUPs result in tissue-specific diseases (Raices and D’Angelo, 2012). Furthermore, in later years many studies have pointed out several transport-independent functions of the NUPs in regulation of genome architecture, stability and organization of peripheral heterochromatin as well as in regulation of gene expression.

In a previous study we identified nucleoporins as important regulators of cyst cell function and cyst cell-germline coordination (Tamirisa et al., 2018). In this study, we go one step further and highlight the critical function of three new nucleoporins in *Drosophila* testis cyst cells, the central channel FG-rich Nup62, the cytoplasmic ring proteins Nup214 and Nup88, and of the unique *Drosophila* Exportin encoded by the *embargoed (emb)* gene. Cyst cells with knockdown of *nup62, emb, nup214* or *nup88* function were not able to encapsulate the germline, became lost over time and could not support germline differentiation. In the absence of cyst cells derived signals, early germ cells overproliferated and testis homeostasis was severely impaired. Suppression of apoptosis could partially restore the defects and reverse cyst cell elimination. Interestingly, overexpression of the *Drosophila* Profilin gene *chickadee (chic)* could rescue cyst cell survival, germline encapsulation, and restore germline differentiation and spermatogenesis. Further analysis indicated that the defects were due to a block in mRNA export from the nucleus, as indicated by nuclear accumulation of the mRNA carrier Rae1. Interestingly, the function Profilin in cyst cells was also associated with mRNA export, independent of Nxt1 and by counteracting most likely the function of Ntf-2, both involved in nucleocytoplasmic transport. This work provides new insights on cell-type specific coordination of nuclear export in *Drosophila* testis cyst cells and how known nucleocytoplasmic regulators namely Nup62, Nup214, Nup88 and Exportin, cooperate with Profilin to support cyst cell function and promote germline differentiation.

## RESULTS

### Nucleoporins and exportin are required in the *Drosophila* testis cyst cells to promote germline encapsulation and differentiation

In an RNAi screen for cyst cell specific regulators in the *Drosophila* testis, we identified that the central channel protein Nup62, the exportin Emb, and the cytoplasmic filament proteins Nup214 and Nup88 [encoded by the *members only (mbo)* gene] control critical steps of spermatogenesis. The *Drosophila* genome contains only one exportin gene, encoded by *embargoed (emb)* gene (also called *crm1* and *xpo1*). Phenotypic analysis and characterization was performed using the *UAS-GAL4* system (Brand and Perrimon, 1993) along with the cyst cell *c587-GAL4* driver (Kai and Spradling, 2004) to knockdown *nup62, emb, nup214* and *nup88* in the *Drosophila* 3^rd^ instar larval (L3) testes. Cyst cell knockdown of any of the four genes in L3 testes resulted in progressive loss of spermatogonia and spermatocyte containing cysts and in non-cell autonomous effects in the germline associated with overproliferation of early germ cells as shown by immunostaining with LamDm0 that marks the nuclear membrane of early germ cells (GSCs, GBs and early spermatogonia) and cyst cells (Fig. 1C-1H). Immunostaining with the nuclear membrane spermatocyte marker LamC revealed that spermatocyte cysts were reduced (Fig. 1J, 1L, 1N, 1P) and spermatocytes were progressively lost (Fig. 1K, 1M, 1O, 1Q) and therefore germline differentiation was significantly impaired. In early steps of the phenotype early germ cell overproliferation was restricted to the anterior part of the testes (Fig. 1J, 1L, 1N, 1P) but as the phenotype advanced resulted in massive filling up of the whole testes with small DAPI-bright germ cells (Fig. 1K, 1M, 1O, 1Q). Staining the same cyst cell knockdowns for Actin showed that the germ cells were not encapsulated in spermatogonial or spermatocyte cysts as in the control testes, and that the branched fusome, present normally in interconnected germ cells, was lost and only dotted spectrosome was observed (Fig. S1A-S1E).

To elucidate whether the overproliferating germ cells were gonialblast-like cells or consisted also of early transit amplifying (TA) spermatogonia, control L3 testes and testes with *nup62* depleted cyst cells were stained for Actin to visualize the presence of spectrosome or fusome and pTyr to assay for the presence of ring canals seen normally in interconnected TA spermatogonia and spermatocytes (Fig.S1F-S1J‘’). Stainings in control testes showed the presence of spherical spectrosome in GSCs and gonialblasts, and of branched fusomes and ring canals in spermatogonial and spermatocyte cysts (Fig.S1F-S1F’’). On the contrary, *c587>nup62^RNAi^* testes showed primarily germ cells with spectrosome and occasionally collapsing TA spermatogonia, which still contained ring canals (Fig.S1G-S1J’’). This observation was consistent with the progression of the phenotype, which showed first loss of spermatogonial cysts and then overproliferation of gonialblast-like germ cells (Fig. 1C-1Q).

Similarly, acute cyst cell knockdown of *nup62, emb, nup214* and *nup88* in adult testes in the background of a *αtubGal80^ts^* (for 4 days and 7 days) showed also overproliferation of early germ cells, stained for Vasa (Fig. 2), suggesting that the function of these genes is consistent in L3 and adult testes. Similar to L3 testes, in 4-days knockdowns overproliferation of early germ cells was restricted to the anterior part of the testes (Fig. 2A-2J) while in 7-days knockdowns the phenotype advanced with massive germ cell overproliferation (Fig. 2K-2T). Expression of a membrane (m)CD8-GFP transgene showed that cyst cells depleted of *nup62, emb, nup214* and *nup88* function were not able to encapsulate the germ cells (Fig. 2C-2J, 2M-2T) in contrast to control adult testes (Fig. 2A, 2B, 2K, 2L). Staining adult testes of the same knockdown genotypes for Actin confirmed the loss of germline encapsulation in spermatogonial or spermatocyte cysts and the presence of only dotted spectrosomes in overproliferating early germ cells (Fig. S2B-S2H‘’) in contrast to the branched fusome in interconnected spermatogonial and spermatocyte cysts of control testes (Fig. S2A-S2A’’). Immunostaining for Traffic-Jam (TJ) (marking CySC and early CCs surrounding spermatogonia) (Li et al., 2003) showed that the majority of TJ-positive cells were lost and few of them surrounded the overproliferating germ cells marked with LamDm0 in 4-days knockdowns or were pushed at the periphery of the anterior testis tip in 7-days knockdowns (Fig. 2 and Fig. S2).

**Figure 2.**
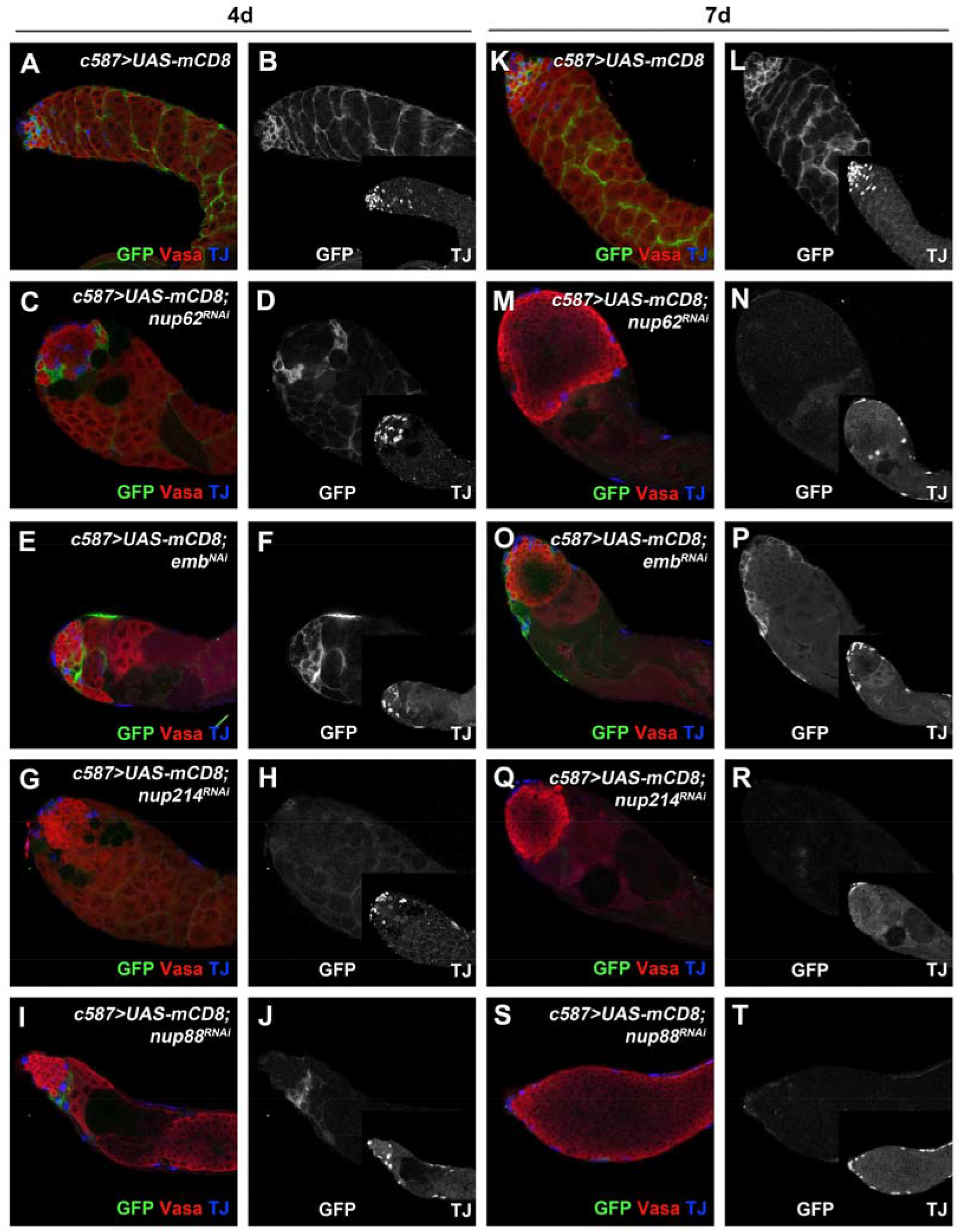
**Knockdown of *nup62, nup214, nup88* and *emb* in cyst cells lead to similar phenotypes in adult as in larval testes**. Adult testes of the indicated genotypes in the background of the *Gal80^ts^* driving expression of a membrane(m)-CD8-GFP (green) in cyst cells co-stained for Vasa (red; germline) and TJ (blue; early cyst cell nuclei). All crosses were performed in the background of the *Gal80^ts^* at 18°C and newly hatched flies of the indicated genotypes were shifted at 30°C to activate the RNAi for 4 days **(A-J)** and 7 days **(K-T)**. Small inset pictures show the TJ-staining only, with TJ-positive cells at the periphery of the overproliferating early germ cells and the anterior testis tip. Testes are oriented anterior left. Image frame: 246μm This figure is associated with **Figures S2** and **S3**.

The inability of *nup62, emb, nup214* and *nup88* depleted cyst cells to properly encapsulate the germline was also observed in L3 *Drosophila* testes with the *UAS-mCD8-GFP* transgene, showing that the membranes of cyst cells could not wrap the germ cells (Fig. 3A-3J). Loss of germ cell encapsulation by the cyst cells was further confirmed by immunostaining for βPS-integrin (Fig. 3K-3O), the β-chain of the integrin heterodimer that marks the cyst cells membranes (Papagiannouli et al., 2014), and Armadillo (Arm; the *Drosophila* homologue of β-Catenin) (Fig. 3P-3T) that marks the hub and the cortical membranes of the squamous cyst cells (Papagiannouli and Mechler, 2009). Importantly, overexpression of *arm* using a *UAS-arm* transgene could not restore germline encapsulation and therefore could also not rescue the observed phenotypes (Fig.S3C). Also, expression of a *UAS-nup214* transgene could not rescue the *nup62* cyst cell knockdown phenotype, suggesting that the two nucleoporins cannot compensate for each other’s function (Fig.S3B).

**Figure 3.**
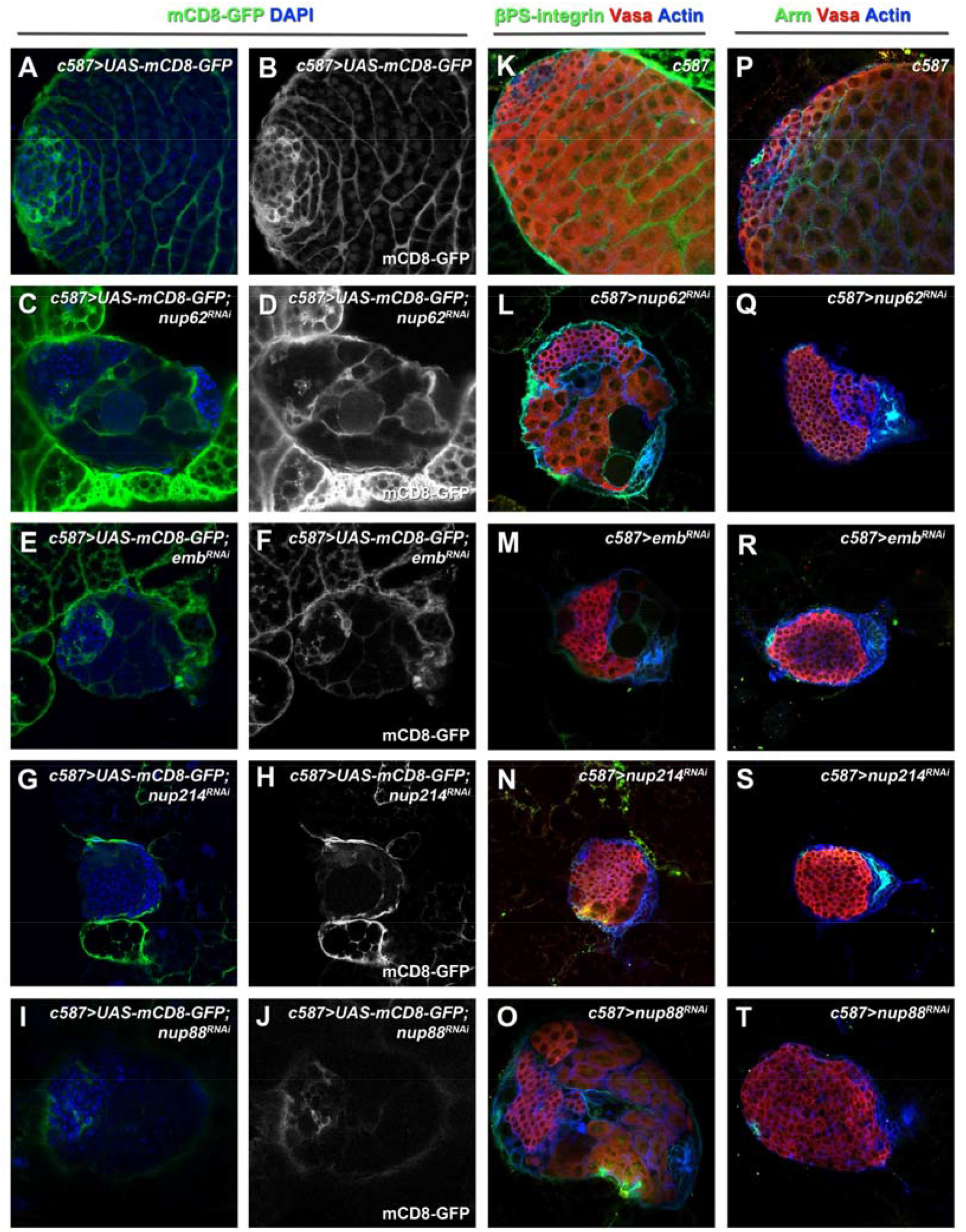
**Knocking down the function of *nup62, nup214, nup88* and *emb* in the cyst cells results in loss of germline encapsulation**. L3 testes of the indicated genotypes **(A-J)** driving expression of mCD8-GFP (green) in the cyst cells co-stained with DAPI (blue) for the DNA, **(K-O)** stained with βPS-Integrin (green) for the cyst cell membranes, Vasa (red) for the germline and DAPI (blue), and **(P-T)** stained with Arm (green) for the cyst cell membranes and the hub, Vasa (red; germline) and DAPI (blue; DNA). Testes are oriented anterior left. Image frame: 246μm

Similar to the adult testes, immunostaining of *nup62, emb* or *nup214* cyst cell knockdowns of L3 testes for TJ and FasIII [normally marking the hub cells of the niche (Patel et al., 1987)], showed that the TJ-positive early cyst cells were getting lost, while FasIII was similar to the wild type testes marking exclusively the hub cells (Fig.S3F-S3K). This result was particularly interesting since previous studies have shown that Piwi and its upstream regulator TJ are required for suppressing FasIII expression in the cyst cells, and loss of *tj* or *piwi* function results in loss of germline encapsulation, ectopic FasIII upregulation in cyst cells and early germline overproliferation (Gonzalez et al., 2015; Li et al., 2003; Saito et al., 2009). Therefore we could conclude that the germline overproliferation we observed in the *nup62, emb* or *nup214* depleted cyst cells was not related to the Piwi-pathway.

Taken together, loss of *nup62, emb, nup214* and *nup88* function in cyst cells resulted in similar cyst cell autonomous as well as germ cell non-autonomous defects in larval and adult testes. Cyst cells could not build and/or maintain cytoplasmic extensions in order to embrace and enclose the germline, while over time cyst cells were completely lost. In the absence of germline encapsulation spermatogonial and spermatocyte cysts collapsed and disappeared, resulting in loss of germline differentiation and overproliferation of early germ cells.

### Suppression of apoptosis in cyst cells can partially restore the defects of *nup62, emb, nup214* and *nup88* knockdowns

Previous studies have shown that overproliferation of early germ cells results non-cell autonomously when cyst cell function is compromised by forced activation of the pro-apoptotic gene *grim*, by impaired germ cell encapsulation e.g. upon loss of the actin polymerization protein Profilin^chic^ or when cyst cells cannot instruct germ cell differentiation due to signaling loss (Ayyub et al., 2015; Dominado et al., 2016; Gonzalez et al., 2015; Li et al., 2003; Qian et al., 2015; Saito et al., 2009; Shields et al., 2014; Tamirisa et al., 2018). Consistently, we found that the viability of the cyst cells was largely restored when apoptosis was prevented in *nup62, emb, nup214* and *nup88* cyst cells knockdowns in L3 and adult testes by co-expression of the anti-apoptotic baculovirus protein p35 (Hay et al., 1994) (Fig. 4 and S4). In testes expressing the *UAS-p35* transgene in the background of *nup62, emb, nup214* or *nup88* cyst cells knockdowns, we observed that TJ-positive cyst cells survived in L3 and adult testes (Fig. 4 and S4). Cyst cells were again able to extent membrane extensions and encapsulate the germ cells, leading to re-appearance of relatively normal spermatogonial and spermatocyte cysts containing branched fusome (Fig. 4F, 4I, 4L, 4O and S4A-S4O). Moreover, no overproliferating germ cells could be observed anymore (Fig. 4D-4O), as LamDm0 staining for early germ cells was restricted at the apical tip of the testes similar to the control testes (Fig. 4A-4C, 4P). However, suppression of apoptosis could only partially restore the developmental defects. The architecture of the rescued spermatogonial cysts was compromised, shown by the abnormal Actin accumulation and thickening of the cyst cell membranes (Fig. S4A-S4L). Moreover, the overall organization of the cysts at the apical testis tip and architecture around the male stem cell niche was abnormal, in comparison to the organized wild type pattern (Fig. 4A-4P and S4M-S4O). However, nuclear localization signal (NLS)-containing proteins could enter the nuclei of cyst cells expressing p35 in the background of *nup62* and *emb* knockdowns (Fig.S4M-S4O). These partially rescued testes were scored as “partially rescued”, since their main defects were reversed to normal.

**Figure 4.**
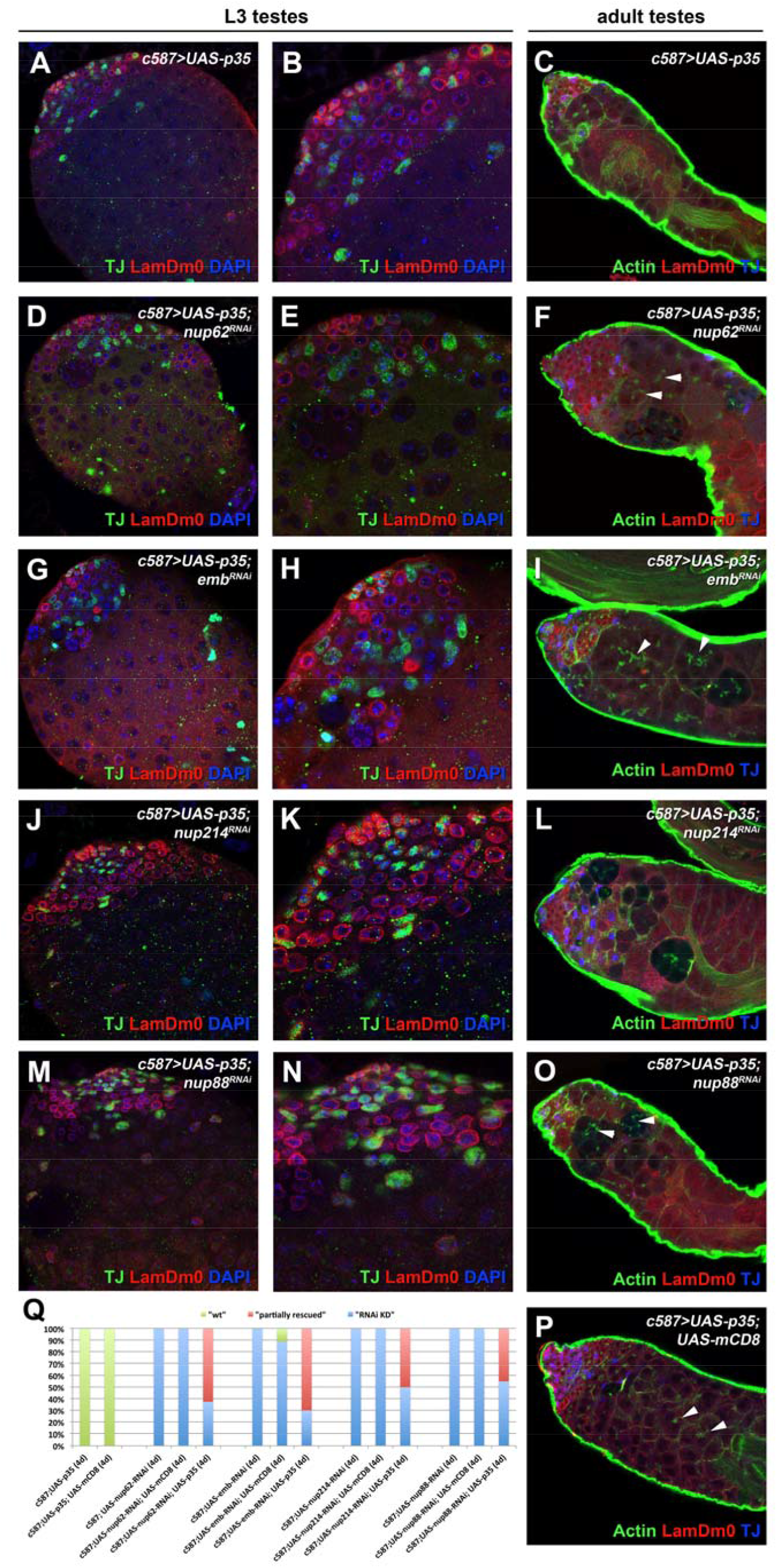
**Suppression of apoptosis in the cyst cells of L3 and adult *Drosophila* testes can restore to a large extent the defects of *nup62, nup214, nup88* and *emb* cyst cell knockdowns**. Testes of control **(A-C)**, *nup62* **(D-F)**, *emb* **(G-I)**, *nup214* **(J-L)**, *nup88* **(M-O)** cyst cell knockdowns expressing a *UAS-p35* transgene to suppress apoptosis, were stained: for TJ (green; early cyst cell nuclei), LamDm0 (red; early germline and cyst cells) and DAPI (blue; DNA) in L3 testes; for Actin (green; phalloidin), LamDm0 (red) and TJ (blue) in adult testes. (B, E, H, K, N) are enlargements of the testis apical region of (A, D, G, J, M). Crosses for adult testes were performed in the background of the *Gal80^ts^*. **(Q)** Quantifications of the different rescue phenotypic classes accompanying each genotype, organized in order of phenotypic strength: weak and strong “RNAi knockdown (KD)”, “partially rescued” and “wt”/“wt rescued” testes. Testes are oriented anterior left. Arrowheads point at branched fusomes. Image frame: (A, B, D, E, G, H, J, K, M, N) 123μm, (C, F, I, L, O, P) 246μm. This figure is associated with **Figure S4**.

The effectiveness of the partial rescue was quantified in adult testes shifted to 30°C for 4 days to activate the transgenes (*UAS-RNAi* and *UAS-p35*) in cyst cells (Fig. 4Q). To control for possible effects of multiple *UAS* constructs, limiting the effectiveness of the *GAL4* driver, control flies carried the same number of *UAS* transgenes using the *UAS-mCD8-GFP* transgene in control genotypes. For example, the rescue strength of flies with the genotype *c587>UAS-nup62^RNAi^; UAS-p35* was compared to those with the genotype *c587>UAS-nup62^RNAi^; UAS-mCD8-GFP*. Interestingly, the rescue showed that the percent of adult testes with *nup62, emb, nup214* or *nup88* cyst cell knockdown phenotype (classified as “RNAi KD”) was substantially reduced usually to less that 50%, as suppression of apoptosis could partially rescue testes to 45-70% (classified as “partially rescued”) from the overall adult testes scored and in comparison to control wild type testes (classified as “wt”).

### Profilin can rescue the defects of *nup62, nup214, nup88* and *emb* knockdowns in cyst cells

Our phenotypes in *nup62, emb, nup214* and *nup88* depleted cyst cells showed striking similarities with the previous described phenotypes of *profilin^chic^* (thereafter mentioned as *profilin*) knockdown in cyst cells (Fig. 5R, S2I, S5C, S5D) (Fairchild et al., 2015; Shields et al., 2014). Profilin, encoded by the *chickadee (chic)* gene in *Drosophila*, is an actin-binding protein involved in a wide-range of actin-based and other cellular processes (Cooley et al., 1992; Theriot and Mitchison; Verheyen and Cooley, 1994) including nuclear export (Minakhina et al., 2005; Stuven et al., 2003). Overexpression of *profilin* in cyst cells, using a *UAS-profilin* transgene (Kulshammer and Uhlirova, 2013) in the background of *nup62, emb, nup214* and *nup88* depleted cyst cells, could perfectly rescue the mutant phenotypes: L3 and adult testes showed no overproliferation of early germ cells, cyst cells survived and could encapsulate the germline giving rise to normal spermatogonial and spermatocyte cysts, germline differentiation as well as testis architecture were perfectly restored (Fig. 5A-5Q, S5E-S5G). Unlike the p35 rescue, Profilin-mediated rescue of *nup62, emb, nup214* and *nup88* cyst cell knockdowns led to rescued testes comparable to the wild type controls (Fig.S2A-S2A’’, 5M, S5A, S5B), which were classified as “wt rescue”.

**Figure 5.**
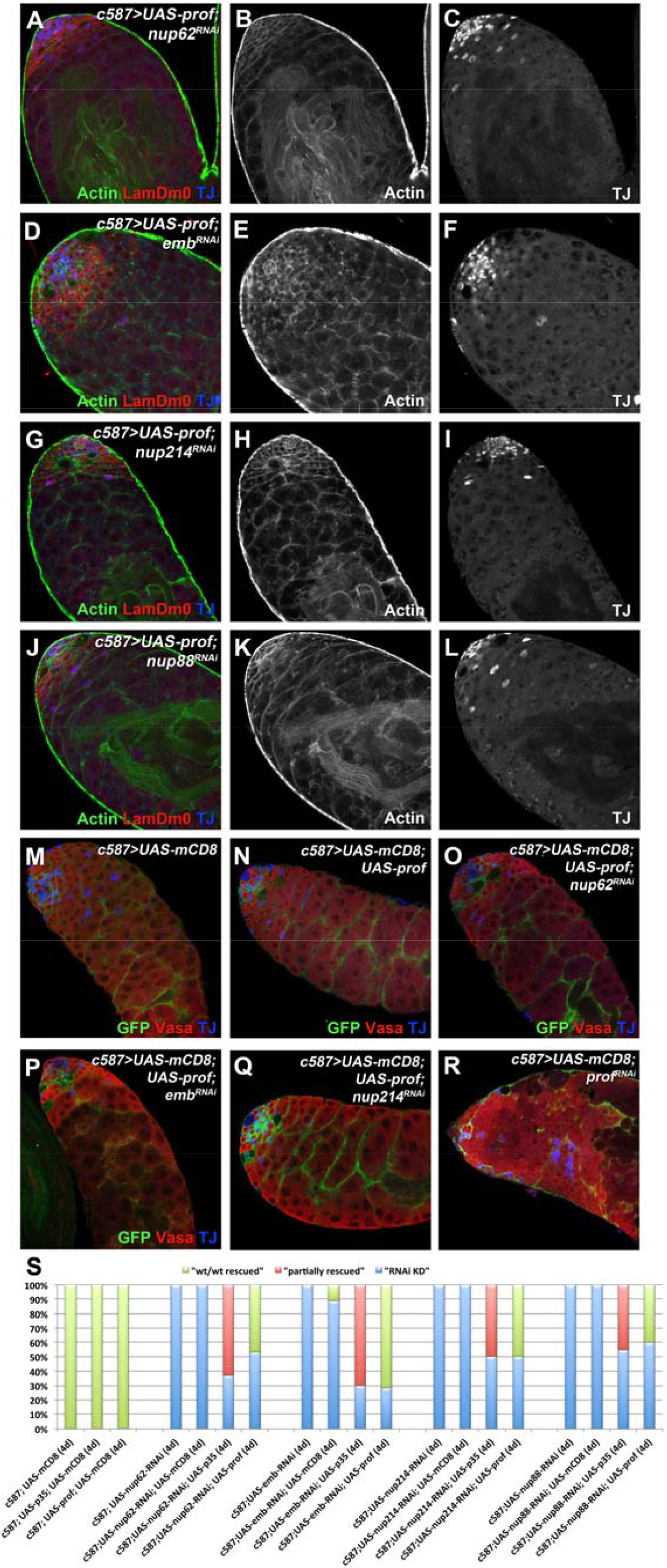
**Expression of *profilin* in cyst cells can fully restore the defects of *nup62, nup214, nup88* and *emb* cyst cell depletion. (A-L)** Adult testes of the indicated genotypes in the background of the *Gal80^ts^* stained for LamDm0 (red; early germline and cyst cells), Actin (green; cyst cells, germline spectrosome and fusome) and TJ (green; early cyst cells) show full rescue of the *nup62, emb, nup214* and *nup88* cyst cell knockdown phenotypes upon co-expression of a *UAS-profilin* transgene. (B, E, H, K) show the Actin staining only and (C, F, I, L) the TJ staining only of (A, D, G, J). **(M-R)** Adult testes of the indicated genotypes marked with mCD8-GFP (green), Vasa (red; germline) and TJ (blue; early cyst cells) show in (O-Q) full rescue of *nup62, emb*, and *nup214* cyst cell knockdown phenotypes upon *profilin* overexpression, in comparison to (N) control flies overexpressing *profilin* and (R) *profilin* cyst cell knockdown with overproliferating early germ cells. mCD8-GFP shows that cyst cells can encapsulate the germline and restore proper architecture in rescued testicular cysts (O-Q) but not in *profilin* depleted cyst cells (R). **(S)** Quantifications of the different rescue phenotypic classes accompanying each genotype, organized in order of phenotypic strength: weak and strong “RNAi knockdown (KD)”, “partially rescued” and “wt”/“wt rescued” testes. Here, rescue efficiency via expression of *p35* is compared to rescue with *profilin*. Newly eclosed male flies were shifted at 30°C to activate the RNAi for 4 days. Testes oriented with anterior at left. Image frame: 246μm. This figure is associated with **Figure S5**.

As with the p35 rescue, the effectiveness of the rescue with profilin was quantified in adult testes shifted to 30°C for 4 days to activate the transgenes (*UAS-RNAi* and *UAS-profilin*) in the cyst cells (Fig. 5S). Control flies carried the same number of *UAS* transgenes using the *UAS-mCD8-GFP* transgene in control genotypes. The percent of adult testes with *nup62, emb, nup214* or *nup88* cyst cell knockdown phenotypes (classified as “RNAi KD”) was substantially reduced to less that 50%, as overexpression of *profilin* gave rise to rescued testes of 45-70% resembling the control wild type (wt) testes (classified as “wt rescued”) from the overall adult testes scored, and in comparison to control testes (classified as “wt”). Interestingly, the percentage of rescued testes in *nup62, emb, nup214* or *nup88* cyst cell knockdowns, either by suppression of apoptosis or by overexpression of *profilin* lead to comparable rescue efficiency e.g. *c587>UAS-emb^RNAi^; UAS-p35* and *c587>UAS-emb^RNAi^; UAS-profilin* (compare in Fig. 5S: the red column “partially rescued” to the green column “wt/wt rescued”). Yet, overexpression p35 not only didn’t rescue the *prof* cyst cell knockdown phenotype (Fig. S6A-S6C) but even more increased the severity of the *profilin* cyst cell depletion phenotype leading to phenotypic enhancement (Fig.S6K).

### Nuclear mRNA export is blocked in cyst cells depleted of *nup62, nup214, nup88, emb* or *profilin* function

Nucleopore proteins of the NPC and exportin allow the selective nucleocytoplasmic transport of macromolecules outside of the nucleus, including mRNA export (Gozalo and Capelson, 2016; Knockenhauer and Schwartz, 2016). To investigate whether loss of *nup62, emb, nup214, nup88* or *profilin* function affects mRNA export in cyst cell nuclei, we assayed for the nuclear vs. cytoplasmic distribution of the mRNA export factor Rae1, by expressing a *UAS-Rae1-GFP* transgene. Rae1 is a shuttling nucleoporin, member of the WD40 protein family, and carrier of mRNA out of the nucleus (Kristo et al., 2017; Sitterlin, 2004). Knockdown of *nup62, emb, nup214, nup88* or *profilin* in cyst cells led to higher Rae1 levels in cyst cell nuclei and strong reduction of Rae1 in the cytoplasm of cyst cells (Fig. 6B-6F’), while control testes exhibited uniform nuclei vs. cytoplasmic Rae1 distribution (Fig. 6A, 6A’). Quantification of fluorescence in cyst cell nuclei (Fig. 6I) further confirmed that Rae1 was retained in the nucleus, and could not shuttle and export mRNA out of cyst cell nuclei in the knockdown phenotypes tested. Notably, cyst cell depleted of *profilin* showed the highest Rae1-GFP accumulation in cyst cell nuclei (Fig. 6I). Quantifications were done with control flies that carried the same number of *UAS* transgenes using the *UAS-mCD8-RFP* transgene in control testes to quantify correctly Rae1-GFP levels in cyst cells e.g. *c587> UAS-Rae1-GFP; UAS-mCD8-RFP* control vs. *c587> UAS-nup62^RNAi^; UAS-Rae1-GFP* flies. Importantly, rescuing *nup62* and *emb* cyst cells knockdowns with p35 (Fig. 6G-6H’) could not restore mRNA export from cyst cell nuclei, since Rae1-GFP levels in cyst cell nuclei remained at the same increased levels as in the corresponding *nup62* and *emb* cyst cells knockdowns (Fig. 6I). This later result indicated that defects in cyst cell survival and germline encapsulation represent most likely secondary effects of *nup62* and *emb* loss that could be partially restored by suppressing apoptosis, while key regulatory functions of cyst cells related to mRNA export remained defective.

**Figure 6.**
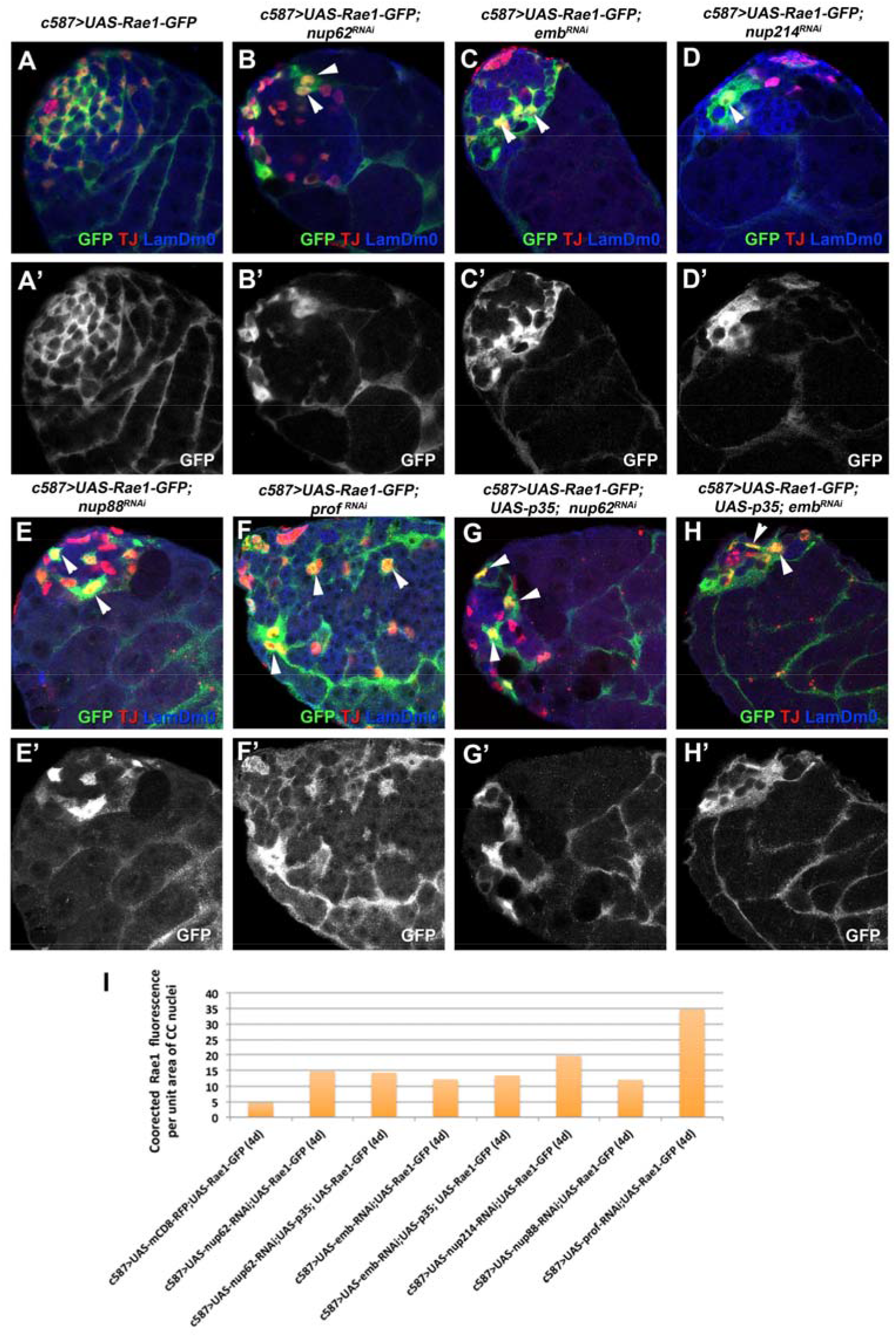
**Knockdown of *nup62, emb, nup214, nup88* and *profilin* in cyst cells leads to nuclear accumulation of the mRNA carrier Rae1. (A-H)** Adult testes of the indicated genotypes in the background of *Gal80^ts^* expressing a *UAS-Rae1* transgene in cyst cells stained for GFP (green), LamDm0 (blue; early germline and cyst cells) and TJ (red; early cyst cell nuclei). Lower panels show the Rae1-GFP staining only. **(I)** Quantification of corrected fluorescent Rae1 levels in cyst cell (CC) nuclei in the indicated genotypes. Arrowheads point at selected Rae1- and TJ-positive cyst cell nuclei. Testes oriented with anterior at left. Image frame: 123μm. This figure is associated with **Figure S7** and **S8**.

### Knockdown of *Nxt1* and *Ntf-2* in cyst cells results in similar phenotypes to loss of *nup62, emb, nup214* and *nup88* function

In a search for proteins that regulate nucleocytoplasmic transport in cyst cells, we investigated the function of the NTF2-related export protein 1 (Nxt1) regulating mRNA export from the nucleus and the Nuclear transport factor-2 (Ntf-2) regulating the nuclear import and export of proteins containing a nuclear localization (NLS) and a nuclear export signal (NES), respectively. Both proteins had also known and predicted interactions with Nup62, Nup214, Nup88 and the exportin Emb analyzed here, based on *STRING (Search Tool for the Retrieval of Interacting Genes/Proteins)* (Fig. 7K). Knocking down *Nxt1* and *Ntf-2* in cyst cells of L3 and adult testes led to phenotypes similar to those of *nup62, emb, nup214* or *nup88* cyst cell knockdowns (Fig. 7A-7F). Interestingly, the block in mRNA export seen in *nup62, emb, nup214* or *nup88* knockdowns was also confirmed for *Nxt1* and *Ntf-2* (Fig. 7G-I), as cyst cells depleted of *Nxt1* and *Ntf-2* function showed elevated nuclear Rae1-GFP levels (Fig. 7J). In order to investigate whether Ntx1 and Ntf-2 act independently or along the same pathway with Nup62, Nup214, Nup88 and Emb mediated mRNA export, thereafter referred to as “Emb-mediated” export, we tested whether overexpression of *profilin* could rescue the *Nxt1* or *Ntf-2* cyst cell knockdown phenotypes, similar to *nup62, emb, nup214* and *nup88*. Interestingly, we obtained a different effect since overexpression of *profilin* in the background of *Nxt1* cyst cell knockdown didn’t influence the phenotypic strength or the penetrance of the phenotype (Fig. S6D-S6F, S6K), suggesting that Profilin acts independent of Nxt1. On the contrary, overexpression of *profilin* in the background of *Ntf-2* depletion in cyst cells resulted in enhancement of the knockdown phenotype. This was manifested in the severity of the phenotype since overexpression of *profilin* led to stronger early germ cell overproliferation covering a larger area in the anterior part of the testes (Fig.S6G-S6I), and in increased penetrance of the knockdown phenotype (Fig.S6K). Presumably, Profilin promotes Emb-mediated export by counteracting the function of Ntf-2 and independent of Nxt1.

**Figure 7.**
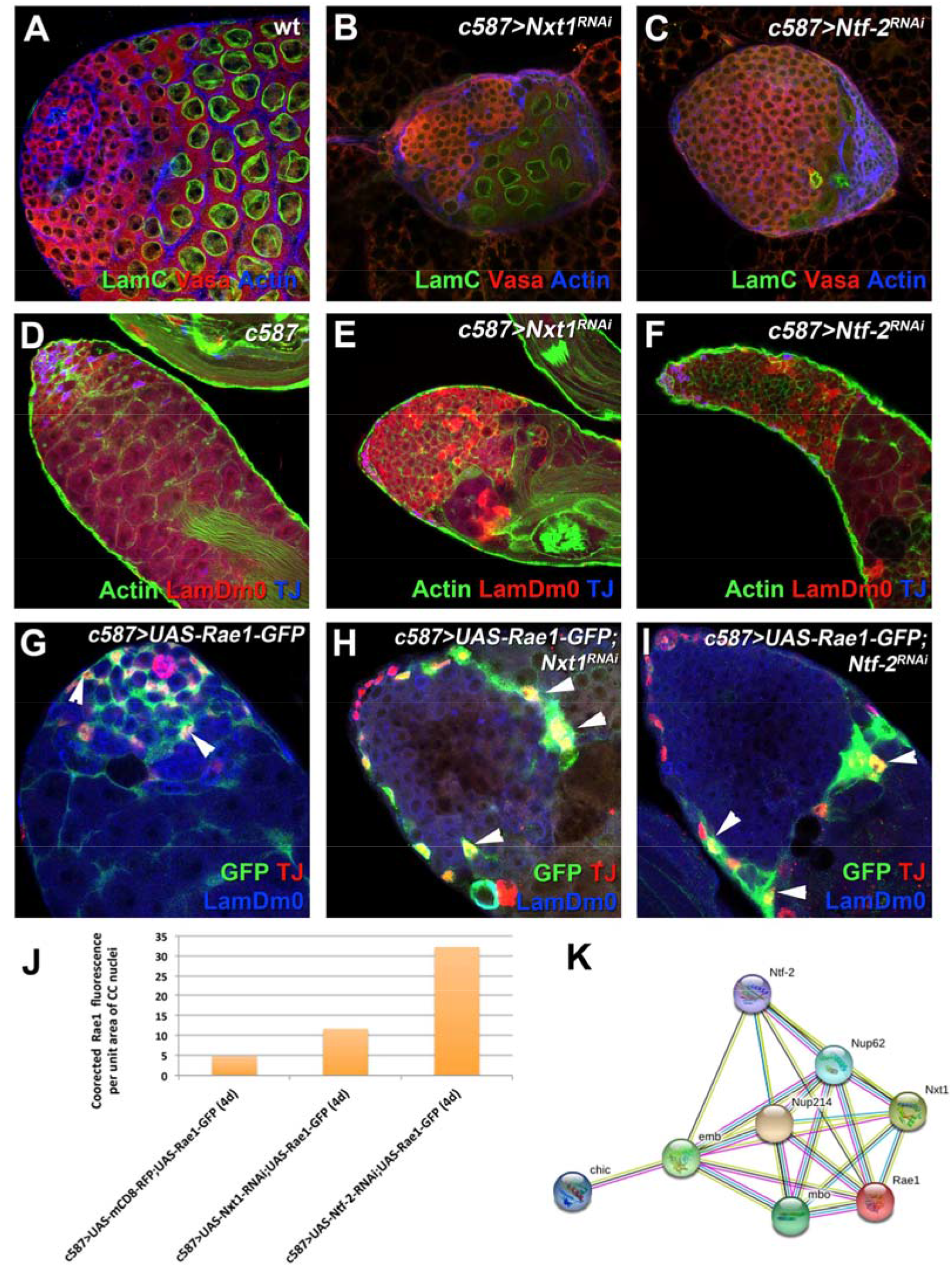
**Nxt1 and Ntf-2 have critical functions in cyst cells associated with defects in mRNA export and non-autonomous early germline overproliferation, similar to loss of *nup62, emb, nup214* and *nup88*. (A-C)** L3 testes of the indicated genotypes stained for LamC (green; spermatocytes), Vasa (red; germline) and TJ (blue; early cyst cell nuclei). Adult testes of the indicated genotypes, in the background of the *Gal80^ts^*: **(D-F)** stained for LamC (green; spermatocytes), Vasa (red; germline) and TJ (blue; early cyst cells); **(G-I)** expressing a *UAS-Rae1* transgene in cyst cells stained for GFP (green), LamDm0 (blue; early germline and cyst cells) and TJ (red; early cyst cell nuclei). **(J)** Quantification of corrected fluorescent Rae1 levels in cyst cell (CC) nuclei in the indicated genotypes. **(K)** Gene network showing interactions of Nup62, Nup214, Nup88 and Emb with Rae1, Profiilin, Nxt1 and Ntf-2, identified in this study (mbo: Nup88, chic: profilin, emb: exportin). Pink: known interactions experimentally determined, Blue: known interactions from curated databases, Black: co-expression associations, Green: text-mining associations. Arrowheads point at selected Rae1 and TJ positive cyst cell nuclei. Testes are oriented anterior left. Image frame: (A-H’) 123μm. This figure is associated with **Figures S6** and **S7**

### Emb and Ntf-2 control the export of NES-containing proteins

As Emb controls export of nuclear export signal (NES)-containing proteins and in many systems NES-mediated protein export depends on mRNA export (Culjkovic-Kraljacic and Borden, 2013; O’Hagan and Ljungman, 2004), we asked whether knockdown of any of components studied so far resulted in defects in protein export. To this end, we used a *UAS-NLS-NES-GFP* transgene, C-terminally tagged with GFP, that contains a wild-type nuclear localization signal (NLS) and nuclear export signal (NES), for which has been previously shown that the activity of the NES is stronger than that of the NLS (Minakhina et al., 2005; Stade et al., 1997). Thus, under normal conditions GFP is observed in the cytoplasm and only when export is impaired GFP remains restricted in the nucleus (Minakhina et al., 2005; Stade et al., 1997). In line with this, in control testes the NLS-NES-GFP localized in the cytoplasm of adult and L3 the cyst cells (Fig.S7A, S8A, S8B), whereas in *nup62, emb, nup214* or *nup88* cyst cell knockdowns the distribution of NLS-NES-GFP was significantly reduced in the cytoplasm of cyst cells (Fig. S7B-S7D, S7I, S8C, S8D). Moreover, we could observe NLS-NES-GFP in cyst cell nuclei depleted of *nup62* and *emb* function (Fig. S7B, S7C; arrowheads), indicating defects in the export of NES-containing proteins, but not in cyst cells depleted of *nup214* or *nup88* (Fig. S7D, S7I). Notably, this effect could be partially restored when *p35* was expressed in the background of *nup62, emb* or *nup214* cyst cell knockdowns, with NLS-NES-GFP decorating again the cytoplasm of cyst cells encapsulating the germ cells, even though the architecture of spermatogonial and spermatocyte cysts at the apical testis tip was not perfectly restored (Fig.S7E-S7H, S8E, S8F). Yet, we could still detect nuclear retention of NLS-NES-GFP in *emb* depleted cyst cells (Fig.S7G; arrowheads), suggesting that in the case of exportin, p35-mediated cyst cell survival could not perfectly restore the export of NES-cargo proteins. Furthermore, NLS-NES-GFP was exported in the cytoplasm of *prof* and *Nxt1* depleted cyst cells, suggesting that they do not regulate the export of NES-containing proteins (Fig.S7J, S7K). Finally, knockdown of *Ntf-2* in cyst cells resulted in accumulation of NLS-NES-GFP in cyst cell nuclei, indicating that Ntf-2 also regulates the export NES-cargo proteins in cyst cells of *Drosophila* testis (Fig.S7L; arrowheads), as in other tissues.

## DISCUSSION

Nucleocytoplasmic communication is of outmost importance for proper cell function, cell survival, signaling response and communication with the neighboring cells and the local tissue microenvironment. While the nuclear membranes form a physical barrier separating the cytoplasm from the nucleus, the nucleopore complex (NPC) forms specialized and dynamic channels that penetrate the nuclear membranes and maintain a selective trafficking of molecular components between the two compartments. As a general rule, the scaffold or transmembrane NUPs are typically stably associated with the NPC and have extremely long turnover rates. On the other hand, peripheral NUPs including many FG-rich NUPs form a meshwork that determines the pore permeability and are in part mobile with dwell times in the minute range (Knockenhauer and Schwartz, 2016; Raices and D’Angelo, 2012; Terry and Wente, 2009). Our work shows how the central channel Nup62, the cytoplasmic Nup214 and Nup88, and the *Drosophila* exportin Emb (also known as Crm1) are critically required in testis cyst cells to maintain cyst cell survival and function in germline encapsulation, to protect cyst cell-germline communication and thereby promote germ cell differentiation. Loss of *nup62, emb, nup214* or *nup88* function via RNAi knockdown in cyst cells, has dramatic effects in spermatogenesis as compromised cyst cell function results in cell non-autonomous overproliferation of early germ cells in the absence of cyst cell derived differentiation signals. Our results have further shown that the cell-autonomous effects in cyst cells depleted of *nup62, emb, nup214, nup88, profilin, Nxt1 and Ntf-2* function are associated with defects in mRNA export, as the mRNA carrier Rae1 accumulates in cyst cell nuclei and cannot shuttle out of the nucleus. Moreover, the exportin Emb and Ntf-2 regulate the export of NES-containing proteins in cyst cells, with Profilin facilitating the Emb- and antagonizing the Ntf-2- mediated nuclear export (Fig. 8).

mRNA export is a multi-step process that involves association with ribonucleoproteins (RNP) to mRNP particles along with several key interactions with the NPC and FG-NUPs (Gozalo and Capelson, 2016; Terry and Wente, 2007). Bulk mRNA export is generally mediated by binding of mRNPs to the Nxf1-Nxt1 complex and other adaptor RNA-binding proteins including Rae1 (Gozalo and Capelson, 2016; Kohler and Hurt, 2007; Okamura et al., 2015). In line with our observations, the exportin Crm1 is also involved in the nuclear export but of a different and smaller subset of specific-mRNAs (including also unspliced or partially spliced mRNAs) (Culjkovic-Kraljacic et al., 2012; Kohler and Hurt, 2007). These mRNAs are connected to the Crm1-dependent export pathway by binding NES-containing adaptor proteins, while efficient export of NES-containing proteins requires ongoing synthesis and export of mRNA (Kohler and Hurt, 2007; O’Hagan and Ljungman, 2004). Yet, both bulk Nxt1-mediated and specific Crm1-mediated mRNA export relay on the FG-rich Nup62 and Nup214 as well as Nup88, which affect the recycling and release steps of mRNAs through the NPC permeability barrier (Culjkovic-Kraljacic and Borden, 2013; Terry and Wente, 2007). As Nxt1 is also implicated in tRNA and in Crm1-dependent export of small nuclear (sn)-RNAs and NES-containing proteins (Zenklusen and Stutz, 2001), and is necessary for the terminal step of Crm1-mediated nuclear export (Black et al., 2001), Nxt1 may participate in both Crm1-dependent and Crm1-independent pathways. Interestingly, *in vitro* and cell-based studies have shown that Nup62 and Nup214 compete for binding Crm1 in mammals albeit Nup214 with a higher affinity, suggesting that the Crm1-export complex can traffic through the NPC by binding progressively to FG-NUPs with higher affinity (Port et al., 2015). Our study confirmed that in the *Drosophila* male germline system mRNA export is mediated by the Nxt1- and as well as the Crm1- export pathways, while Crm1 is also needed for the export of NES-containing proteins. Yet, genetic interaction experiments with overexpression of profilin suggest that Nxt1 probably acts independently of the Exportin/Emb export pathway in testis cyst cells. Therefore, impaired mRNA export is identified as the main cell-autonomous defect underlying the loss of function phenotypes of all the players analyzed in this study. Importantly, suppression of apoptosis can restore only the secondary effects associated with cyst cell survival and germline encapsulation and not the mRNA export defects.

**Figure 8.**
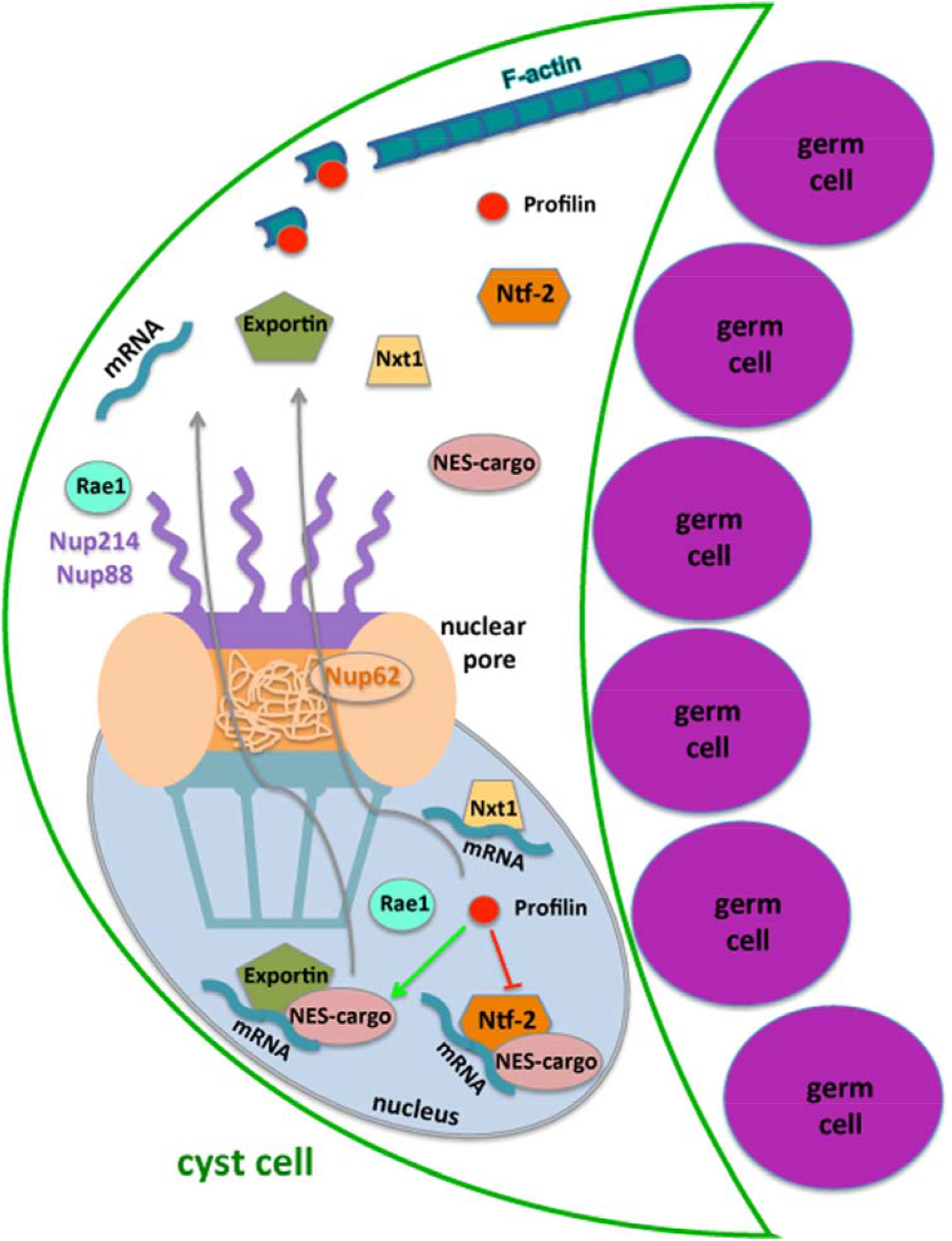
**Model diagram showing how Nup62, Nup214, Nup88 and Exportin channel the export of mRNAs and NES-cargo proteins and the possible involvement of Rae1, Profilin, Nxt1 and Ntf-2**. mRNA is exported from the nucleus of *Drosophila* testis cyst cells in an Exportin (Emb/Crm1)-mediated way along with NES-containing proteins or in a bulk Nxt1-dependent way. Ntf-2 also affects export of mRNA and NES-proteins from cyst cell nuclei. Block in mRNA export results in Rae1 retention in cyst cell nuclei. Profilin facilitates Emb-mediated export and antagonizes Ntf-2-mediated export through a yet unknown mechanism, with no effect on Nxt1, while in the cytoplasm Profilin regulates actin polymerization.

The involvement of Nup62, Nup214 and Nup88 in the export of NES-containing proteins is not very clear. In cyst cells depleted of *nup62*, NLS-NES-GFP is lost from the cytoplasm but we can observe nuclei with NLS-NES-GFP content while p35 expression restores NLS-NES-GFP in the cytoplasm and NLS-GFP in nuclei of cyst cell suggesting that both import and export function properly. In *nup214* and *nup88* cyst cell knockdowns, NLS-NES-GFP is lost from both the cytoplasm and nuclei of cyst cells, while p35 rescue restores the NLS-NES-GFP in the cytoplasm. One possible explanation is that NLS-NES-GFP is exported in the cytoplasm but as cyst cells lose their cytoplasmic extensions, it gets progressively lost. Alternatively, a block in nuclear import upon loss of *nup214* and *nup88* prevents NLS-NES-GFP from entering the cyst cell nuclei. Previous studies in *Drosophila* have shown that Nup88 anchors CRM1 at the nuclear envelope and down-regulates the levels of NES protein export, while NLS import or mRNA export are not affected (Roth et al., 2003). This suggests that the role of these proteins in nucleocytoplasmic transport may vary depending on the functional requirements of a particular cell-type. Overall, we cannot conclude from our data to what extend the defects in mRNA export could be coupled to the defects in export of NES-proteins in *Drosophila* testis cyst cells, as in other systems. However, we have shown that Profilin can influence the function of the two main players involved in NES-protein export by supporting the function of Emb and antagonizing the function of Ntf-2.

Another effect of loss of *nup62, emb, nup214* or *nup88* function is that cyst cells are progressively eliminated, an effect that could be rescued to a large extend by expression of the pro-apoptotic gene p35. Similar to our observations, deregulation of Nup214 in myeloid precursor cells leads to nuclear mRNA accumulation and apoptosis (Boer et al., 1998). In mammalian cells, mutations in the NES domain of the adaptor protein Crk can abolish binding to Crm1 and promote apoptotic death in these cells (Smith et al., 2002). Therefore, block in the export of mRNA or NES-cargo proteins is many cases associated with apoptosis, across different tissues. Nowadays it is well established that components involved in mRNA export, NES-protein export and the NPC barrier and/or transport function, are very often implicated in proliferation, survival, signaling or metastasis. Even more, NES-cargo proteins that bind Crm1 in order to be transported out of the nucleus can be tumor suppressors, oncoproteins or even pro-apoptotic genes (e.g. p53, BRCA1, Survivin, APC), which normally regulate transcription, chromosomal architecture, alternative pre-mRNA splicing or cell cycle control (Nguyen et al., 2012; Schwerk and Schulze-Osthoff, 2005). Thus, an imbalance in the cytosolic vs. nucleolar levels can have profound effects in the function of these proteins: e.g. Survivin regulates cell division in the nucleus, while in the cytoplasm acts as cytoprotective by actively inhibiting apoptotic pathways (Musch et al., 2002; Nguyen et al., 2012). Along this line, it is possible that testis cyst cells experiencing mixed, stage inappropriate or apoptotic signals, as a result of the abnormal distribution of regulatory components, become eliminated. On the other hand, mRNA biogenesis involves quality controls that detect possible errors in transcription, mRNA processing, formation of export-competent mRNPs or mRNA export (Kohler and Hurt, 2007; Wickramasinghe and Venkitaraman, 2016). Accordingly, as mRNA export is coupled to transcription, mRNA splicing and maturation, blocking mRNA export could activate distinct levels of post-transcriptional control and DNA damage recognition (Culjkovic-Kraljacic and Borden, 2013; Wickramasinghe and Venkitaraman, 2016), leading to elimination of the cyst cells. Finally, work in *Schizosaccharomyces pombe* revealed that the mRNA export and cell cycle regulator Rae1 has a DNA-damage checkpoint function and gets recruited to double-strand breaks along with Cdc25 and Checkpoint kinase 1 (Chk1) (Selvanathan et al., 2010). Thus in addition to its role in mRNA export, Rae1 may also possess a checkpoint function in testis cyst cells and accumulation of Rae1 in cyst cell nuclei, upon block of mRNA export, may result in activation of DNA damage responses in cyst cells depleted of *nup62, emb, nup214* or *nup88* function. Interestingly, expression of p35 could rescue the survival of cyst cells that could again grow and encapsulate the germ cells but the mRNA export was not restored showing that mRNA export relies on the proper function of these proteins.

Profilin is an actin-binding protein and highly conserved regulator of actin-dependent processes across different species (Cooley et al., 1992; Theriot and Mitchison; Verheyen and Cooley, 1994). It is also a multitasking protein that binds a plethora of diverse ligands, with functions ranging from endocytosis and membrane trafficking to Rac-Rho signaling and nuclear activities (Witke, 2004). In particular, Profilin can enter the nucleus and regulate the export of NES-cargo proteins in *Drosophila* by counteracting the function of Ntf-2 (Minakhina et al., 2005), and facilitate the nuclear export of Profilin-Actin complexes by Exportin-6 in HeLa cells (Stuven et al., 2003). Moreover, Profilin can associate with transcriptionally active genes, co-localize with snRNP-core proteins, Cajal bodies and nuclear structures involved in pre-mRNA splicing (Birbach, 2008; Percipalle, 2013; Skare et al., 2003; Soderberg et al., 2012). Our data in the *Drosophila* testis show that Profilin is required for the export of mRNA from cyst cell nuclei and can rescue the defects in cyst cells introduced upon loss of *nup62, emb, nup214* or *nup88* function. Overexpression of profilin in the background of *nup62, emb, nup214* or *nup88* cyst cell knockdowns, restores cyst cell survival, germline encapsulation and differentiation, and reestablishes normal spermatogenesis. Similar to previous reports in *Drosophila* (Minakhina et al., 2005), Profilin antagonizes the function Ntf-2 in cyst cells as overexpression enhances the defects associated Ntf-2 depletion in cyst cells. As the primary cause underlying the compromised cyst cell function of Profilin, Nup62, Emb, Nup214 and Nup88 relies on defects in nuclear export of mRNA, one can hypothesize that the function of Profilin in cyst cells is tightly linked to Emb-mediated nuclear export by counteracting the function of Ntf-2. The fact that suppression of apoptosis cannot reverse any of the profilin-associated defects, indicates that the key regulatory functions of profilin in cyst cells are not only linked to nuclear export but also to the organization of the actin cytoskeleton in the cytoplasm.

More recent studies in mammalian systems and in *Drosophila* uncovered the role of NUPs in functions beyond nucleocytoplasmic transport, in regulation of chromatin architecture and compartmentation, stability and organization of peripheral heterochromatin and in regulation of gene expression (Hou and Corces, 2010; Kalverda and Fornerod, 2010; Kalverda et al., 2010; Raices and D’Angelo, 2012). Interestingly, in the *Drosophila* ovaries Nup93 and Nup154 recruit the FG-containing Nup62 and together regulate chromatin distribution in nuclei of both somatic and germline cells (Breuer and Ohkura, 2015). In *Drosophila* embryonic cells, Nup62, Nup50 and Nup98 interact with transcriptionally active genes inside the nucleoplasm to stimulate the expression of developmental and cell cycle genes (Kalverda and Fornerod, 2010). Importantly, NUPs can also bind promoters, enhancers and insulators as Nup98 mediates enhancer-promoter looping at ecdyson-inducible genes in *Drosophila*, while Nup93 and Nup153 bind to super-enhancers regulating cell-type specific genes in humans (Ibarra et al., 2016; Pascual-Garcia et al., 2017). Although Nup62 is necessary for nuclear export in cyst cells, it is still plausible that this function may be coupled to transport-independent functions in gene regulation and chromatin compartmentation in cyst cells.

Furthermore, the protein composition of NUPs can vary among different cell types and tissues with distinct developmental roles, and this explains why mutations in different NUPs result in tissue-specific defects and diseases (Raices and D’Angelo, 2012). In the *Drosophila* testis, Nup154 is required for spermatogonial maintenance and *nup154* mutants are linked to increased cell death and reduced cell proliferation by deregulating the Dpp pathway (Colozza et al., 2011). Loss of Nup98-96 leads to dramatic cell autonomous effects in *Drosophila* male germ cells, since germ cells differentiate prematurely to spermatocytes at the 2-, 4-, and 8- germ cell cysts stage (Parrott et al., 2011). Moreover, loss of *nup205, nup93-1, nup358, nup44A* in *Drosophila* testis cyst cells leads to an overproliferation of early germ cells (Tamirisa et al., 2018). Finally, LamDm0 acts downstream of the EGFR pathway and together with the nuclear basket Nup153 and Mtor regulate nuclear retention of the phosphorylated MAPK dpERK (Chen et al., 2013).

Our study highlights a cell-type specific requirement of distinct peripheral nucleoporins, exportin and profilin in nuclear export in *Drosophila* testis cyst cells. Block in mRNA export is toxic for the cyst cells demonstrating the importance of nucleocytoplasmic transport for cyst cells survival and function in supporting the developmental decisions of the germline and in testis homeostasis.

## EXPERIMENTAL PROCEDURES

### Fly stocks and husbandry

*Oregon R* was used as a wild type stock. The following stocks were obtained from the Bloomington Stock Center (BL) Indiana: *UAS-nup62-RNAi^TRiP.HMC03668^, UAS-nup214-RNAi^TRiP.HMS00837^, UAS-nxt1-RNAi^TRiP.GL00414^, UAS-Ntf-2-RNAi^TRiP.JF03048^, UAS-NLS-NES-GFP* (BL7032), *Pin/CyO; UAS-mCD8-GFP* (BL5130), *UAS-p35* (BL5072, BL5073), *UAS-mCD8-GFP* (BL5139). The following stocks used in this study were obtained from the Vienna *Drosophila* RNAi Center (VDRC) Austria: *UAS-nup62-RNAi^v100588^, UAS-nup88-RNAi^v47692^, UAS-emb-RNAi^v103767^ UAS-profilin-RNAi^v102759^*. *UAS-profilin^chic^/CyO* was a gift of Mirka Uhlrova and *UAS-Rae1-GFP* a gift of Peter Vilmos and Chunlai Wu (Tian et al., 2011). Other fly stocks used in this study are described in FlyBase (www.flybase.org). All *UAS-gene^RNAi^* stocks are referred to in the text as *gene^RNAi^* for simplicity reasons. Knockdowns were performed using the *UAS-GAL4* system (Brand and Perrimon, 1993) by combining the *UAS-RNAi* fly lines with cell-type specific *GAL4* drivers described above.

For the phenotypic analysis in 3^rd^ instar larval (L3) testes: (a) *c587-GAL4* flies were crossed to *UAS-gene^RNAi^* flies and crosses were raised at 30°C (these gave rise to strong phenotypes), or (b) *c587-GAL4> Gal80^ts^* flies were crossed to *UAS-gene^RNAi^* flies, crosses were raised at 18°C and were shifted at 30°C as 1^st^ instar larvae for two days and were dissected at L3 (these gave rise to weaker phenotypes). For the phenotypic analysis in adult testes, *c587-GAL4>Gal80^ts^* flies (in combination with other *UAS* transgenic flies used in this study) were crossed to *UAS-gene^RNAi^*, crosses were raised at 18°C until male adults hatched. Then males with the correct genotype (along with few females) were shifted at 30°C for 4 days or 7 days and then phenotypes were analyzed.

### Immunofluorescence staining and microscopy

Whole mount testes immunostained were dissected in PBS, fixed for 20min in 8% formaldehyde, rinsed in 1% PBX (1% Triton-100x in PBS) and blocked in 5% Bovine Serum Albumin in 1% PBX. Testes were incubated with primary antibodies overnight at 4°C and the following day with the secondary antibodies for 2h at room temperature in the dark (Papagiannouli et al., 2014). For testes immunostaining in the presence of GFP, 1% PBT (1% Tween-20 in PBS) was used instead of 1% PBX in all steps. Testes were mounted in ProLong^®^ Gold Antifade (Thermo Fischer Scientific). The monoclonal antibodies used in this study: anti-Armadillo N7A1 (1/10; mouse), anti-Vasa (1/10; rat), anti-βPS-integrin-CF.6G11 (1/10; mouse), anti-LamC-LC28.26 (1/10; mouse), anti-LamDm0-ADL101 (1/10; mouse), anti-FasIII-7G10 (1/10; mouse) were obtained from the Developmental Studies Hybridoma Bank developed under the auspices of the NICHD and maintained by The University of Iowa, Department of Biological Sciences, Iowa City, IA 52242. Polyclonal anti-Vasa-dC13 (goat; 1/20) was from Santa Cruz Biotechnology. Monoclonal anti-Phosphotyrosine Antibody clone 4G10 (mouse; 1/200) was from Millipore, chicken anti-GFP (13970; 1/10,000) from Abcam and guinea-pig anti-TJ (1:5000) polyclonal antibodies was a gift of Dorothea Godt. Filamentous Actin (F-actin) was stained with Alexa Fluor phalloidin 488, 546 or 647 (1/300, Thermo Fischer Scientific) and DNA with DAPI (Thermo Fischer Scientific). Following secondary antibodies were used: donkey anti-goat Alexa Fluor-488, donkey anti-mouse Alexa Fluor-546 and Alexa Fluor-647, donkey anti-guinea pig Alexa Fluor-546 and Alexa Fluor-647, donkey anti-rat Alexa Fluor-488, donkey anti-rabbit Alexa Fluor-488 and Alexa Fluor-546 from Thermo Fischer Scientific (1/500) and donkey anti-chicken Alexa Fluor-488 from Jackson ImmunoResearch (1/500).

Confocal images were obtained using a Leica system SP8 (1024×1024pix, 184μm image frame) (Beckmann Center “Cell Sciences Imaging Facility”, Stanford University; COS, University of Heidelberg). Pictures were finally processed with Adobe Photoshop 7.0.

## AUTHOR CONTRIBUTIONS

F.P. conceived, designed, performed, interpreted experiments and wrote the paper. M.T.F. supported the study financially (NIH grant 1-R01 GM080501). I.L. assisted in interpreting experiments and obtained funding to support the study (DFG/SFB 873).

## ACKNOWLEDGMENTS

We would like to thank the *Drosophila* community for providing us generously with fly stocks and antibodies, in particular Dorothea Godt, Mirka Uhlirova, Peter Vilmos, Chunlai Wu, the Developmental Studies Hybridoma Bank (Iowa State University), the Vienna *Drosophila* Resource Center (VDRC) and the Bloomington *Drosophila* Stock Center. We apologize to all whose work was not sited due to space limitations. We would like to deeply thank Maria Leptin, as part of this work was done in her lab. This work was supported by the DFG/SFB 873 to IL, NIH grant 1-R01 GM080501 to MTF and PA2659/4-1 and PA2659/5-1 to FP. We thank the Cell Sciences Imaging Facility (Beckmann Center, Stanford University) and the CECAD Imaging Facility (University of Cologne) for their technical support.

## Abbreviations

CySCs: somatic cyst stem cells
CC: somatic cyst cells
FG: phenylalanine-glycine
GB: gonialblast
GSCs: germline stem cells
L3: 3^rd^ instar larval stage
NUP: nucleoporin
NPC: nuclear pore complex

